# Developmental IL7/IL7R signaling in fetal resident tissue macrophages regulates adaptive responses to liver injury

**DOI:** 10.64898/2026.10.26.761961

**Authors:** Nicole A. Tseng, Gabriel A. Leung, Brian P. Krum, Ernest M. Kameni, Clint H. Valencia, Samuel Arda, Daniel S. Barrera, Noah Huerta, Patrice N. Mimche, Anna E. Beaudin

## Abstract

Resident tissue macrophages (RTMs) originate from multiple fetal hematopoietic waves and seed organs during the prenatal window where they acquire tissue-specific homeostatic functions. Several growth factors within the fetal niche are required for RTM development, but knowledge of how these signals program tissue-specific RTM survival, identity, and function during the prenatal window is limited. Here, we investigated the functional role of developmental IL7/IL7R signaling on fetal RTM survival and function. IL7 deficiency impaired neonatal RTM establishment, similar to IL7R deficiency, and IL7 was broadly expressed in fetal tissues and co-localized with IL7R+ macrophages. IL7/IL7R signaling regulated cell cycle, apoptotic, and inflammatory transcriptional pathways and activated phospho-AKT signaling in fetal liver myeloid cells. IL7R deletion impaired proliferation of fetal liver monocyte precursors and increased fetal liver macrophage apoptosis. Specific Cre-mediated IL7R deletion in fetal definitive HSC-derived, but not EMP-derived, macrophages caused irrecoverable postnatal loss of IL7R-expressing macrophages in specific tissues, including the liver, demonstrating a specific temporal requirement for IL7R expression during fetal myelopoiesis. Prenatal IL7R blockade subsequently impaired adaptive responses to chronic fibrotic liver injury in adulthood, associated with increased monocyte infiltration and upregulation of pro-inflammatory signaling pathways. These findings redefine IL7 as a developmental myeloid niche factor and demonstrate that IL7/IL7R signaling is co-opted during fetal development to imprint durable RTM programs that shape postnatal tissue immunity and adaptive responses to injury.

## Introduction

Resident tissue macrophages (RTMs) are specialized macrophages embedded within organs, where they execute functions critical for maintaining tissue homeostasis^1^. Genetic fate-mapping approaches have demonstrated a fetal origin for many RTMs, with primitive yolk sac (YS) progenitors, early and late YS erythromyeloid progenitors (EMPs) and definitive hematopoietic stem cells (HSCs) contributing to developmental heterogeneity among some, but not all RTM populations ^2–4^. Both the development of fetal RTMs from pre-macrophages (pMacs) that directly colonize embryonic tissues ^5^, and from fetal monocytes that seed tissues across development ^3^, have been described. Although multiple Cre-based fate-mapping models have been used to study resident tissue macrophage (RTM) origins, the relative contributions of distinct fetal hematopoietic progenitors to specific RTM populations remain unclear. Moreover, how RTM functional diversity is established from different developmental sources during fetal life is unknown.

We previously reported that expression of the lymphoid-associated interleukin-7 receptor (IL7R) regulates RTM development^6^. Originally identified as a bone marrow growth factor inducing proliferation of B cell precursors, interleukin-7 (IL7) is broadly required for lymphoid development ^7–9^. In adult hematopoiesis, the common lymphoid progenitor is the first progenitor in the hematopoietic hierarchy to express IL7R and respond to IL7 signaling, and fate-mapping has demonstrated that IL7R expression is restricted to the lymphoid lineage in adult murine hematopoiesis ^10,11^. IL7 signaling is known almost exclusively as a regulator of lymphocyte proliferation, survival, and differentiation ^7,10,12,13^. In contrast to the lymphoid-restricted role of IL7R signaling in adult hematopoiesis, we demonstrated that IL7R is expressed by fetal myeloid cells and that deletion or prenatal blockade of IL7R impairs cellularity of neonatal RTM compartments ^6^. However, the specific role of the cognate ligand, IL7, and the mechanisms by which IL7/IL7R signaling regulate fetal myeloid developmental pathways are unknown. Macrophage development universally requires M-CSF ^14–19^, but acquisition of RTM identity is also regulated by tissue– and niche-specific factors ^18,20–22,23–27^. We hypothesize that IL7/IL7R signaling is co-opted by developing myeloid cells as an additional signal that regulates specific developmental functions during fetal myelopoiesis.

Here we examined how IL7/IL7R signaling regulates developmental functions during fetal myelopopiesis, including the role of the IL7 ligand, downstream signaling mechanisms, and the developmental requirement of IL7/IL7R signaling for acquisition of postnatal RTM function. We show that genetic loss of IL7 also reduces neonatal RTM compartments, and that IL7 is expressed in the liver and across peripheral tissues in association with IL7R+ macrophages. We also show that IL7/IL7R stimulation in fetal myeloid cells activates pI3K/AKT signaling and regulates macrophage proliferation, survival, and acquisition of an anti-inflammatory phenotype. Cre-mediated deletion of Il7r from different waves of fetal myelopoiesis identified a wave of definitive fetal blockade of IL7R impairs adaptive responses to chronic fibrotic liver injury, associated with alterations to myeloid populations and pro-inflammatory gene expression. Together, our findings support a role for IL7 as an additional developmental myeloid niche factor that supports acquisition of fetal RTM function and regulation of postnatal tissue function.

## Materials and Methods

### Animals

All animals were bred and housed in the UC Merced or University of Utah AAALAC accredited vivaria and all animal procedures were approved by UC Merced and University of Utah Institutional Animal Care and Use Committees (IACUC). All animals were used in experiments regardless of sex, and animals were sacrificed by CO_2_ asphyxiation, and cervical dislocated or decapitated as secondary euthanasia. Mouse lines included C57BL/6 (00664JAX), *IL7^GFP^*provided by the Ikuta Lab ^28^, *Il7r-/-*(002295JAX), Csf1r Mer-icre-Mer (019098JAX), Cdh5 Cre-ERT2 (Taconic #13073), IL7R^fl^ (022143JAX), and Ai9 (007909JAX).

### 4-OHT administration

For induction of Cre activation in the offspring of Csf1r Mer-iCre-Mer crosses, a single 75 mg/kg dose of 4-hydroxytamoxifen (4-OHT) was administered to pregnant dams intraperitonially at E8.5. For the Cdh5Cre-ERT2 crosses, a single 50 mg/kg dose of 4-OHT was administered to pregnant dams intraperitonially at E10.5. 4-OHT solutions were supplemented with 37.5mg/kg or 25 mg/kg of progesterone to reduce adverse effects of tamoxifen on pregnancies. Pups were delivered by cesarean section and cross fostered.

### Tissue processing

Tissue digestion was performed with 0.2-2 mg/mL collagenase IV (Gibco) and 4.8 mg/mL Dispase (Gibco) at 37°C. Single cell suspension was attained by homogenization with 16G and 19G needles, or p1000 pipette and filtered through a 70 µM filter. Total cellularity was determined by bead count in each sample and extrapolated based on bead recovery and divided by tissue weight to determine cells/gram tissue.

### Immunofluorescence Imaging

Embryos were cryosectioned and fixed in 4% formaldehyde and blocked with 1x PBS + 0.1% Triton-X + 10% NGS overnight at 4°C. Primary antibody staining was performed overnight, and secondary antibodies were stained for 1 hour and stained with DAPI. Images were collected on a Zeiss LSM 700 Zen Black software and 63x objective. Image analysis was performed with ImageJ.

### Flow cytometry

Live cells were stained with antibodies in 1x PBS + 5 mM EDTA + 2% serum. For phosphoflow analysis, cells were fixed in 1.5% formaldehyde and either permeabilized with ice cold 100% methanol or permeabilized using the BioLegend True Nuclear Kit (Cat # 424401). Samples were analyzed using the BD LSR II, BD Aria III, or Cytek Aurora. All antibodies used are listed in Supplementary Table 1. Definitions of hematopoietic populations are listed in Supplementary Table 2.

### Gene Expression Analysis

12,000-15,000 CD11b^lo^F4/80^hi^ and CD11b^hi^F4/80^lo^ cells were sorted from WT and *Il7r^-/-^*fetal tissues for sequencing, with the exception of fetal brain F4/80^lo^ cells (4,000-8,000 cells). Most populations were sorted from whole tissue, but brain myeloid cells were enriched using CD11b microbeads (Miltenyi). Definitions of sorted populations are listed in Supplementary Table 2. RNA was isolated from four biological replicates of sorted F4/80^hi^ and F4/80^lo^ cells from *Il7r^-/-^* and WT brain, liver, lung, and skin (Qiagen 74104), and libraries were prepared (NEB E7760L) and sequenced using the Illumina NovaSeq6000, to a depth of 20 million reads. Transcripts were counted using FeatureCounts and differential expression was determined based on an FDR<0.05 using DESeq2 in the Bioconductor RNA-seq workflow.

### Prenatal anti-IL7R blockade

Pregnant C57BL6/J mice received intraperitoneal (i.p.) injections of 600 µg of anti-IL7R blocking antibody (clone A7R34) or an isotype antibody control (IgG2a) at embryonic days (E)15.5 and 17.5 (proliferation/apoptosis assay or (E)13.5, 15.5 and 17.5 (liver fibrosis assays).

### Carbon tetrachloride (CCl_4_) model of hepatic injury and fibrosis

Six week-old female C57BL6/J offspring of mice that had received either control IgG2a or A7R43 IL7R blocking antibody were IP injected with 2 μL/g of 10% CCl_4_ (Sigma-Aldrich) diluted in olive oil three times a week for 6 weeks. Control mice received olive oil only. Mice were euthanized 48h after the final injection of CCl_4_ and the livers and serum collected for histology, flow cytometry, biochemical and molecular analysis.

### RNA extraction, cDNA synthesis and qPCR

Total liver RNA extraction and cDNA synthesis were performed as previously described^29,30^. Real-time qPCR reactions were performed using Quantitect SYBR Green PCR reagent (Qiagen) and qPCR reaction plates run in the Applied BioSystems QuantStudio^TM^ 6 Flex real-time PCR system (ThermoFisher). Primer sequences are described in Supplementary Table 3. Transcripts were normalized to the housekeeping gene *hypoxanthine-guanine phosphoribosyl transferase* (*HPRT*) and mRNA expression levels were calculated using the 2^-ΔΔCt^ method.

### Histology

Livers were fixed in 10% buffered formalin and multiple paraffin-embedded 6µm sections were stained with H&E and Masson’s Trichrome according to standard protocols. High power images were acquired using an EVOS FL Widefield microscope (ThermoFisher).

### Liver damage enzyme detection

Plasma samples from oil and CCl_4_-treated mice were processed in a single batch for determination of serum alanine aminotransferase (ALT) and aspartate aminotransferase (AST) levels using a DC Element chemistry analyzer (HESKA).

### Ex vivo culture

E14.5 fetal monocyte dendritic progenitor (MDP:Monocyte-Dendritic Cell Progenitors (MDPs;cKit+CSF1R+CD11b+Flt3^+^), fetal cMOPs (Ly6c^hi^cKit+CSF1R+CD11b+Flt3-), and fetal myeloid progenitors (MPs; (Ly6c^lo^cKit+CSF1R+CD11b+Flt3-) were sorted from *IL7R^-/-^* and WT fetal livers and cultured for 6 days in Dulbecco’s Modified Eagle’s Media (DMEM) supplemented with 20% bovine calf serum and M-CSF (20 ng/ml, Peprotech) ± rIL7 (100 ng/ml, Peprotech).

### Statistical Analysis

Statistical analysis was performed in Graphpad Prism or DESeq2. Significance of *in vivo* cellularity, frequency, and MFI analysis was determined using the Mann-Whitney test (unpaired, two-tailed) and significance of *ex vivo* cell culture cellularity was determined by unpaired student’s t-test or paired student’s t-test, as indicated.

### Data Sharing Statement

The RNA-sequencing data set is accessible NCBI’s Gene Expression Omnibus through GEO accession number GSE226299.

## Results

### IL7 signaling regulates fetal RTM development

We previously demonstrated that blockade or deletion of the IL7R receptor impairs fetal RTM development. As it is possible for other signals to regulate RTM development through the IL7R receptor, we investigated the role of the cognate ligand for IL7R, IL7, in RTM development. We determined the impact of IL7 deficiency on RTM development in *Il7*-deficient *Il7^GFP/GFP^* knock-in mice and WT controls. Lymphocytes were greatly reduced in *Il7^GFP/GFP^* neonates and adults, as previously reported ^31–33^ **(SFig. 1A-C)**. RTMs in the liver (**Fig. 1A**), brain (**Fig. 1C**), and epidermis (**Fig. 1D**) but not in the lung (**Fig. 1B**) were significantly reduced by *Il7* deficiency at postnatal day (P) 4, similar to what we previously observed with *Il7r* deficiency or fetal IL7R blockade ^6^(**SFig. 2A**). By adulthood, *Il7-*deficient RTM populations recovered to WT levels **(Fig. 1E-H, SFig 2B)**, similar to *Il7r*^-/-^mice ^6^, suggesting that in the absence of IL7/IL7R signaling, RTMs either undergo local proliferation to restore homeostasis or are replenished by alternative sources during adulthood ^6,34,35^. Using an anti-GFP antibody in *IL7^GFP/+^* embryos ^28^, we identified broad IL7 expression in the liver (**Fig. 1I)** and other peripheral tissues at embryonic day (E) 14.5 (**SFig. 1D-F**). IL7R-expressing RTMs identified by co-staining of F4/80 and IL7R localized in clusters in proximity to IL7+ cells, suggesting co-localization. Together, these data suggest that IL7 ligand, and IL7 signaling through IL-7R, regulates early fetal macrophage development.

**Figure 1.**
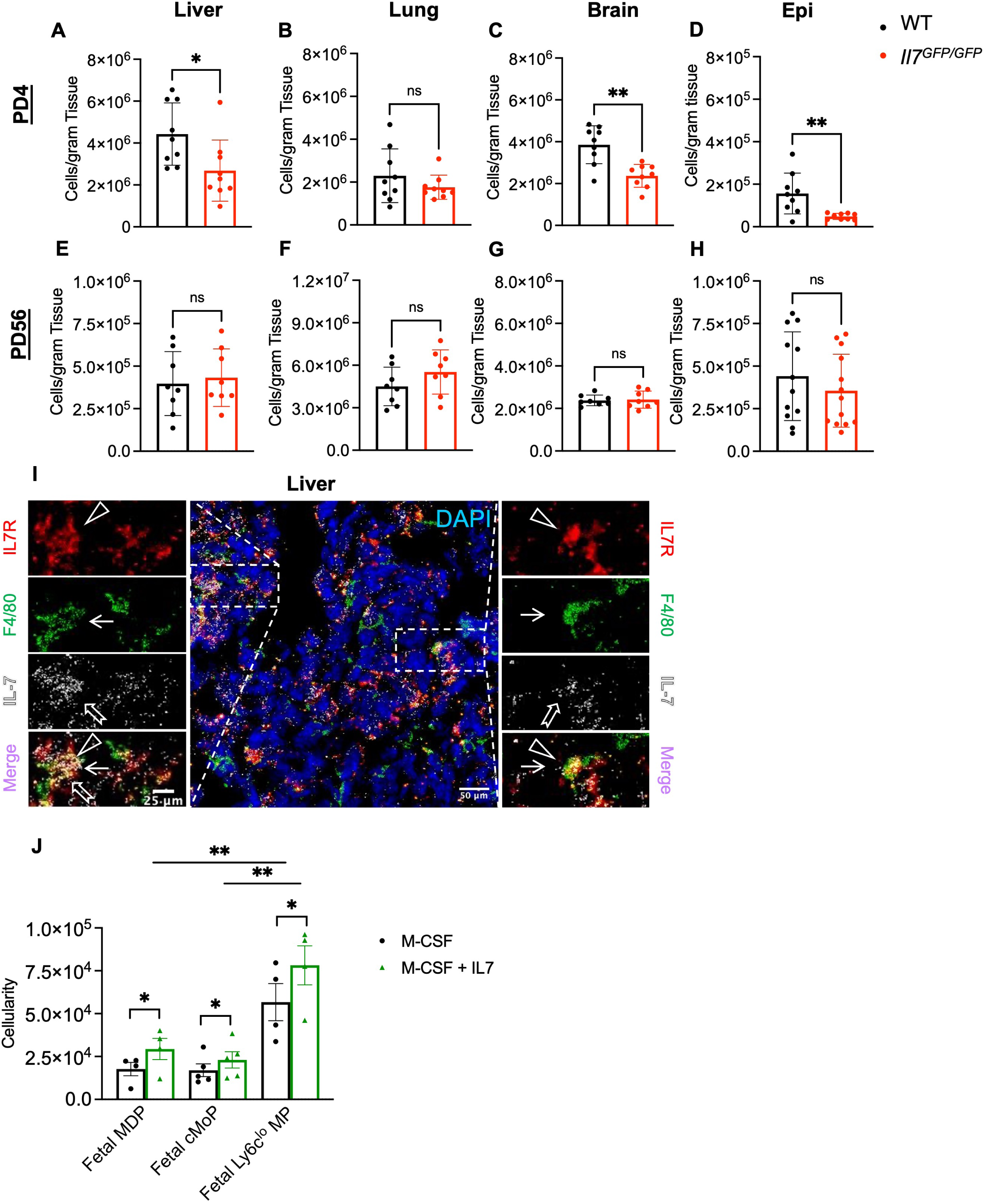
IL7 signaling regulates fetal RTM development. (**A-D**) IL7-deficient Il7^GFP/GFP^ neonatal RTM cellularity compared to WT across tissues. Data represent average ± SD for n=9 mice per condition representing three independent experiments. (**E-H)** *Il7^def^* adult RTM cellularity compared to WT across tissues. Data represents average ± SD of n=6-12 mice per condition representing four independent experiments. Significance determined using Mann-Whitney test, *\*p<0.05, **p<0.01.* **(I**) Immunofluorescence staining of IL7R+ F4/80+ cells in close proximity to IL7 expression in liver. Slides were imaged on a Zeiss LSM 700 Zen Black software and 63x objective, images generated using ImageJ software. Rectangular fields show images at magnification and highlight IL7R+ F4/80+ cells colocalized with IL7+ staining. IL7R (red) indicated by arrowhead, F4/80 (green) indicated by arrow, and IL7 (gray) indicated by fishtail arrow. Images are representative of three individual embryos. **(J)** Cellularity of F4/80^hi^ macrophages generated after 6 days of culture of E14.5 WT fetal MDPs, fetal cMOPs, and fetal MPs in M-CSF alone or M-CSF (20 ng/ml) + rIL7(100 ng/ml). Data represents average ± SD from three individual experiments and progenitors plated in triplicate. Significance determined by paired t-test of averaged triplicate values, *\*p<0.05, **p<0.01, ***p<0.001*.

**Figure 2.**
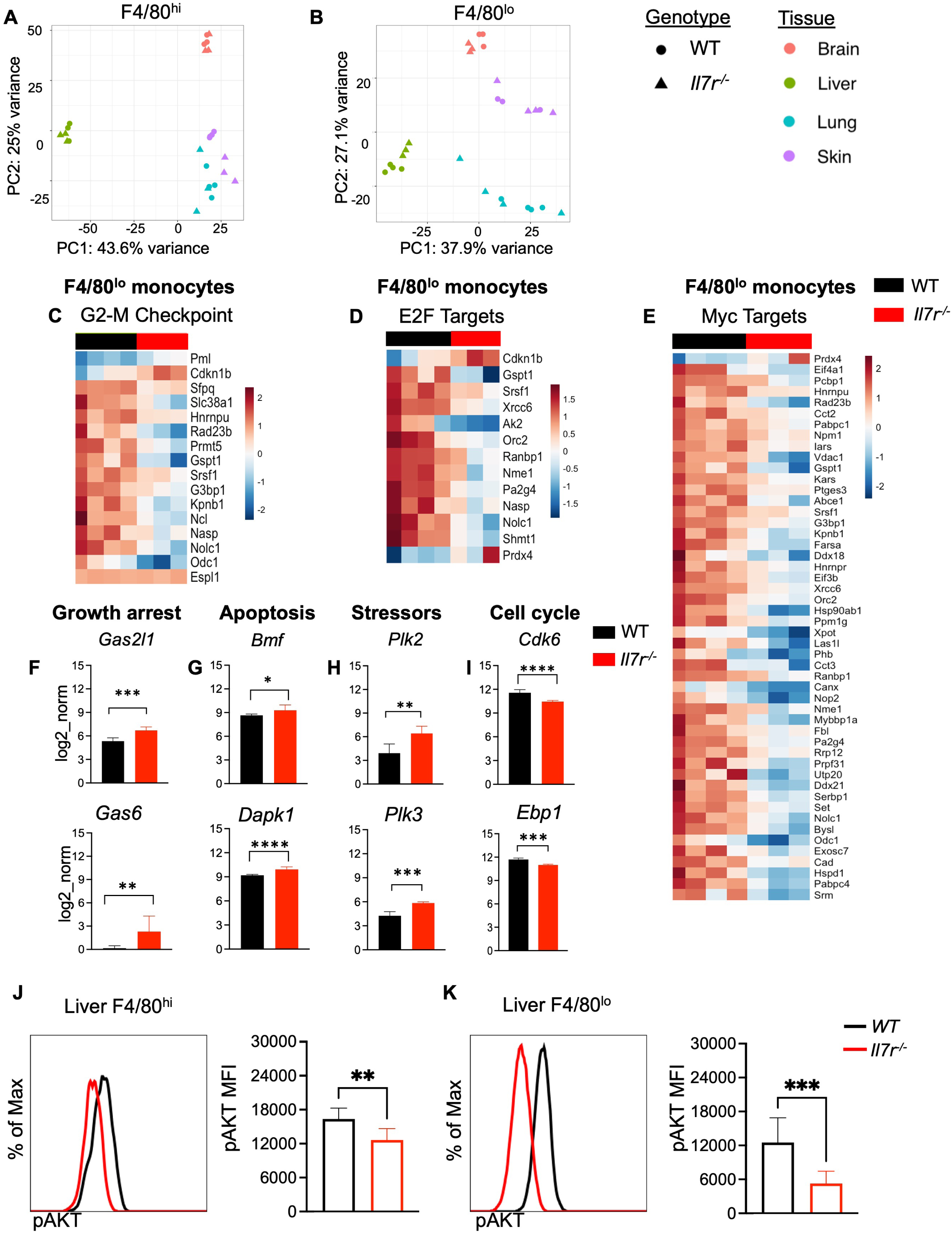
IL7R expression regulates developmental cell cycle pathways and pAKT signaling. Unsupervised clustering of *Il7r^-/-^* and WT F4/80^hi^ (**A**) and F4/80^lo^ cells **(B)** (see **SFig3A** for gating strategy). **(C-E**) Differential expression in liver F4/80^lo^ monocytes within pathways associated with the G2-M Checkpoint **(C)**, the E2F targets **(D**), and Myc Targets **(E). (F-I**) Log_2_ normalized fold change of differential expression of specific genes associated with growth arrest **(F**), apoptosis **(G**), cellular stress **(H**), and proliferation (**I**). Gene expression analysis based on four biological replicates, log_2_ normalized fold change based on FDR < 0.05. (**J,K**)Representative histograms and quantification of of phospho-AKT detection in CD11b^lo^F4/80^hi^ **(J)** and CD11b^hi^F4/80^lo^ cells **(K)** isolated from *Il7r^-/-^* (red line) fetal liver as compared to *WT* mice (black line). Data represent average ± SD of n=3-12 mice per condition for 4 independent experiments. Significance determined using Mann-Whitney test, *\*p<0.05, **p<0.01, ***p<0.001*.

We next tested the ability of IL7 to directly modulate macrophage output from cultured fetal liver (FL) myeloid progenitors *ex vivo*. We tested three previously identified fetal myeloid progenitors: fetal Monocyte-Dendritic Cell Progenitors (MDPs;cKit+CSF1R+CD11b+Flt3^+^), fetal cMOPs (Ly6c^hi^cKit+CSF1R+CD11b+Flt3-), and fetal myeloid progenitors (MPs; (Ly6c^lo^cKit+CSF1R+CD11b+Flt3-), where fetal MPs have been proposed to be EMP-derived and fetal MDPs are proposed to be HSC-derived, with fetal cMOPs as a common intermediate^3^. All three progenitors were cultured with M-CSF ± IL7 and macrophage differentiation was quantified *ex vivo* on day 6 of culture. *Ex vivo* culture with IL7 alone in the absence of M-CSF did not generate significant numbers of macrophages (**SFig. 2C**) indicating that IL7 was not sufficient for macrophage differentiation, similar to TGF-β ^36^. However, the addition of IL7 to M-CSF culture significantly promoted macrophage generation on day 6 from all three progenitor subsets relative to M-CSF alone (**Fig. 1J**), with the greatest macrophage production observed from fetal MPs. Together, *ex vivo* culture experiments demonstrate that while IL7 is not required for macrophage differentiation, IL7 enhances macrophage generation from fetal myeloid progenitors.

### *IL7R* expression regulates developmental cell cycle pathways

To identify downstream transcriptional programs regulated by IL7R signaling, we performed gene expression analysis on E17.5 WT and *Il7r* ^-/-^myeloid cells isolated by differential expression of CD11b and F4/80 in the brain, lung, and skin, as well as Ly6c in the liver (**SFig. 3A**). Unsupervised clustering of sorted populations revealed strong tissue-specific signatures (**Fig. 2A, B**), with genotype-driven clustering emerging only in select populations, including skin F4/80^hi^ cells, brain F4/80^lo^ cells, and liver F4/80^lo^ cells. We have previously shown that FL F4/80^lo^ monocytes initiate Il7r transcription prior to circulation and further upregulate Il7r upon entering the bloodstream ^6^. We therefore hypothesized that Il7r-dependent transcriptional programs first emerge in FL F4/80^lo^ monocytes and subsequently shape the development and establishment of tissue-resident F4/80^hi^ macrophage populations. Accordingly, we focused our analysis on transcriptional differences between *Il7r* ^-/-^ and WT FL F4/80^lo^ monocytes.

**Figure 3.**
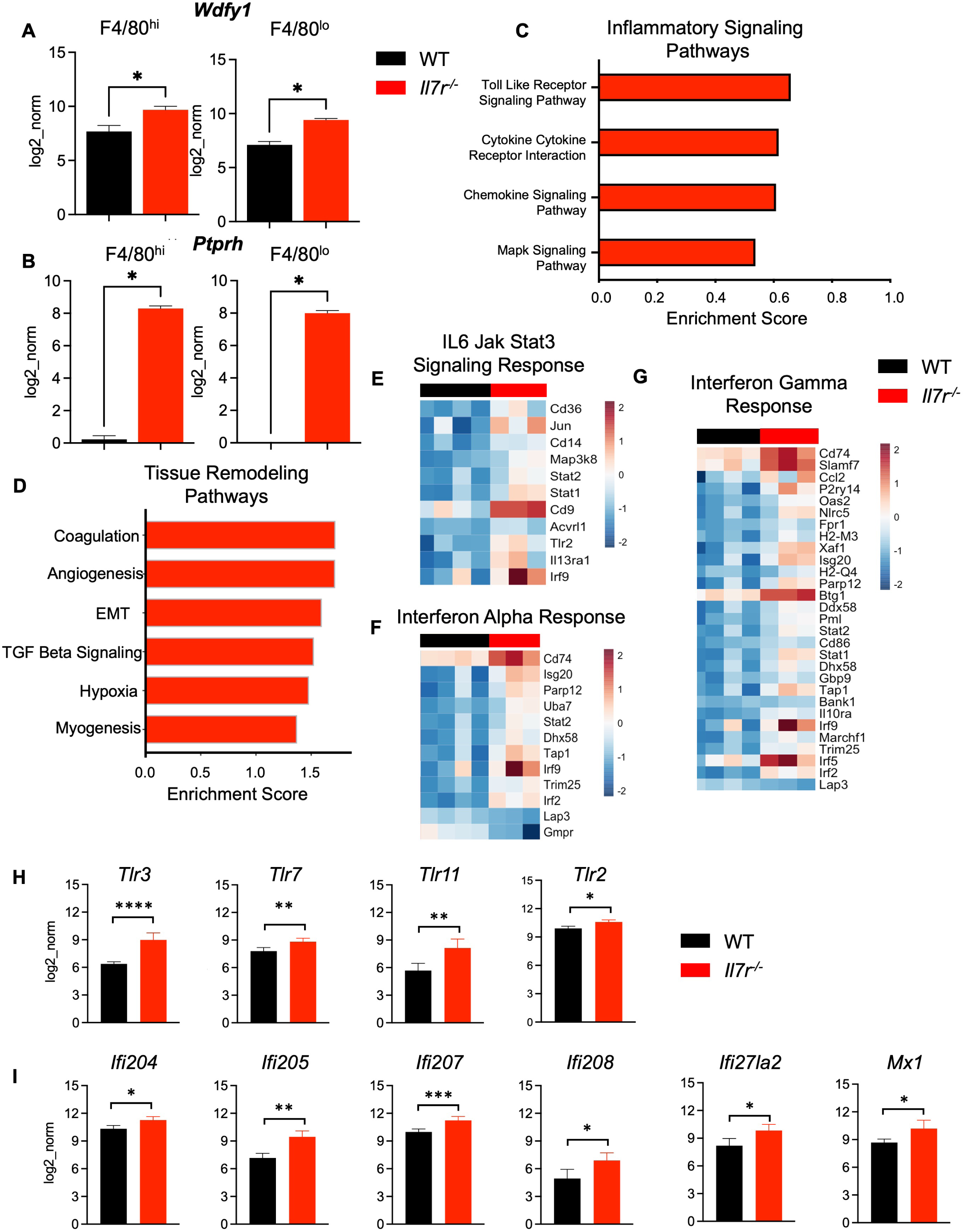
IL7R expression regulates acquisition of an inflammatory transcriptional signature in fetal myeloid cells. (A,B) Top differentially expressed genes in F4/80^hi^ and F4/80^lo^ myeloid cells across all tissues. (**C-D**) Signaling pathways enriched in *Il7r*^-/-^ liver CD11b^hi^F4/80^lo^ cells compared to WT. **(E-G)** Differential expression in fetal liver F4/80^lo^ monocytes of the IL6-Jak-Stat3 signaling pathway (**E**), the interferon alpha response (**F**), and the interferon gamma response pathways **(G**)**. (H-I)** Log_2_ normalized fold change of differential expression in fetal liver F4/80^lo^ cells of Toll-like receptor genes (**H**) and interferon response genes (**I**). Gene expression analysis based on four biological replicates, log_2_ normalized fold change based on FDR < 0.05.

Cell cycle pathways were significantly downregulated in *Il7r* ^-/-^FL Ly6C^hi^F4/80^lo^ monocytes, including G2-M checkpoint pathway (**Fig. 2C**), E2F targets (**Fig. 2D**), and Myc targets (**Fig. 2E**). *Il7r ^-/-^*FL F4/80^lo^ cells specifically upregulated expression of the growth arrest genes *Gas2l1* and *Gas6* (**Fig. 2F**), the apoptotic genes *Bmf* and *Dapk1*, **(Fig. 2G)**, and the polo-like kinases *Plk2* and *Plk3* **(Fig. 2H**), cellular sensors of stress and regulators of the cell cycle ^37,38^. Additionally, *Il7r* ^-/-^FL F4/80^lo^ monocytes also downregulated expression of the proliferation genes, *Cdk6* and *Ebp1* **(Fig. 2I)**, further demonstrating cell cycle dysregulation. These transcriptomic analyses suggested that *Il7r* deficiency impaired cell cycle regulation, proliferation, and survival pathways in F4/80^lo^ FL monocytes.

We next investigated signaling pathways activated by IL7/IL7R signaling in FL myeloid cells to regulate cellular functions, including proliferation and survival. In lymphoid cells, IL7 binding to IL7R rapidly triggers phosphorylation and licensing of Janus Kinase 1 and Janus Kinase 3 (JAK1/3), leading to Stat5 phosphorylation ^39–41^. Stat5 phosphorylation was not detectable in fetal myeloid cells in response to IL7 stimulation (**SFig. 3B,C**). However, *ex vivo* stimulation with GM-CSF for 5 minutes resulted in Stat5 activation in F4/80^lo^ monocytes, as previously reported ^42^, but did not stimulate F4/80^hi^ macrophages isolated from the fetal liver (**SFig. 3D,E**). The addition of IL7 also did not increase Stat5 signal in F4/80^lo^ monocytes in combination with GM-CSF (**SFig. 3F,G**). Thus, IL7 does not signal through Stat5 in fetal myeloid cells.

PI3K-AKT signaling is another prominent pathway downstream of IL7R signaling that promotes lymphocyte cell growth, proliferation, and survival. We compared AKT phosphorylation in freshly isolated E17.5 *ll7r*^-/-^ and WT FL myeloid populations using phospho-flow cytometry without *ex vivo* stimulation (**Fig. 2J,K**). AKT phosphorylation was significantly decreased in both *Il7r* ^-/-^F4/80^hi^ and F4/80^lo^ fetal liver cells as compared to WT **(Fig. 2J,K)**. These data demonstrate that IL7R signaling activates PI3K-AKT signaling in fetal liver myeloid cells – both monocytes and macrophages.

### *Il7r* expression regulates acquisition of an anti-inflammatory transcriptional signature in fetal myeloid cells

In addition to cell cycle and survival pathways, inflammatory signaling pathways were also significantly upregulated in the fetal *Il7r* deficient myeloid transcriptome. The most broadly upregulated genes across all cell types and tissues were *Wdfy1*, an endosomal mediator of toll-like receptor (TLR) 3 signaling (**Fig. 3A**), and *Ptprh*, a protein tyrosine phosphatase (**Fig. 3B**). Within FL monocytes, the TLR signaling pathway was the most upregulated pathway in *Il7r* ^-/-^ liver F4/80^lo^ monocytes **(Fig. 3C)**, but cytokine signaling pathways were also broadly enriched. *Il7r* ^-/-^ FL monocytes also exhibited an upregulation of tissue-remodeling pathways including TGF-β signaling, hypoxia, angiogenesis, and epithelial-mesenchymal-transition pathways, among others (**Fig. 3D**). IL6-Jak-Stat3 signaling (**Fig. 3E**), interferon alpha response (**Fig. 3F**), and interferon gamma response pathways (**Fig. 3G**) were all upregulated in response to *Il7r* deficiency. Multiple TLR genes were upregulated in *Il7r* ^-/-^ FL F4/80^lo^ monocytes **(Fig. 3H)**. Additionally, multiple interferon-inducible genes were upregulated **(Fig. 3I)**, indicating that *Il7r* expression regulated interferon-mediated inflammatory responses and tissue-remodeling pathways in FL monocytes.

### IL7/IL7R regulates proliferation and survival in fetal macrophages

To define the cellular processes governed by IL7/IL7R signaling during RTM development, we assessed proliferation and apoptosis of fetal RTMs at E18.5 using Ki67 and Annexin V staining. For these experiments, we employed an orthogonal approach of temporal IL7R blockade— injecting pregnant C57BL/6 mice with anti-IL7R antibody (A7R34) or isotype control at E15.5 and E17.5 (**Fig. 4A**)—to specifically target the developmental window when fetal myeloid cells express IL7R^6^. Twenty-four hours after blockade, proliferation of F4/80^lo^ monocytes in the fetal liver was modestly reduced (**Fig. 4B,D**), whereas F4/80^hi^ macrophage proliferation was unchanged (**Fig. 4B,C**). In contrast, IL7R blockade markedly increased apoptosis in F4/80^hi^ macrophages and, to a lesser extent, in F4/80^lo^ monocytes (**Fig. 4E-G**). We additionally observed elevated apoptosis in lung myeloid cells and reduced proliferation in brain myeloid populations (**SFig. 4A–B**), indicating tissue-specific responses. Collectively, these data show that IL7R signaling is required to support both survival and proliferation of developing myeloid cells in a tissue-dependent manner.

**Figure 4.**
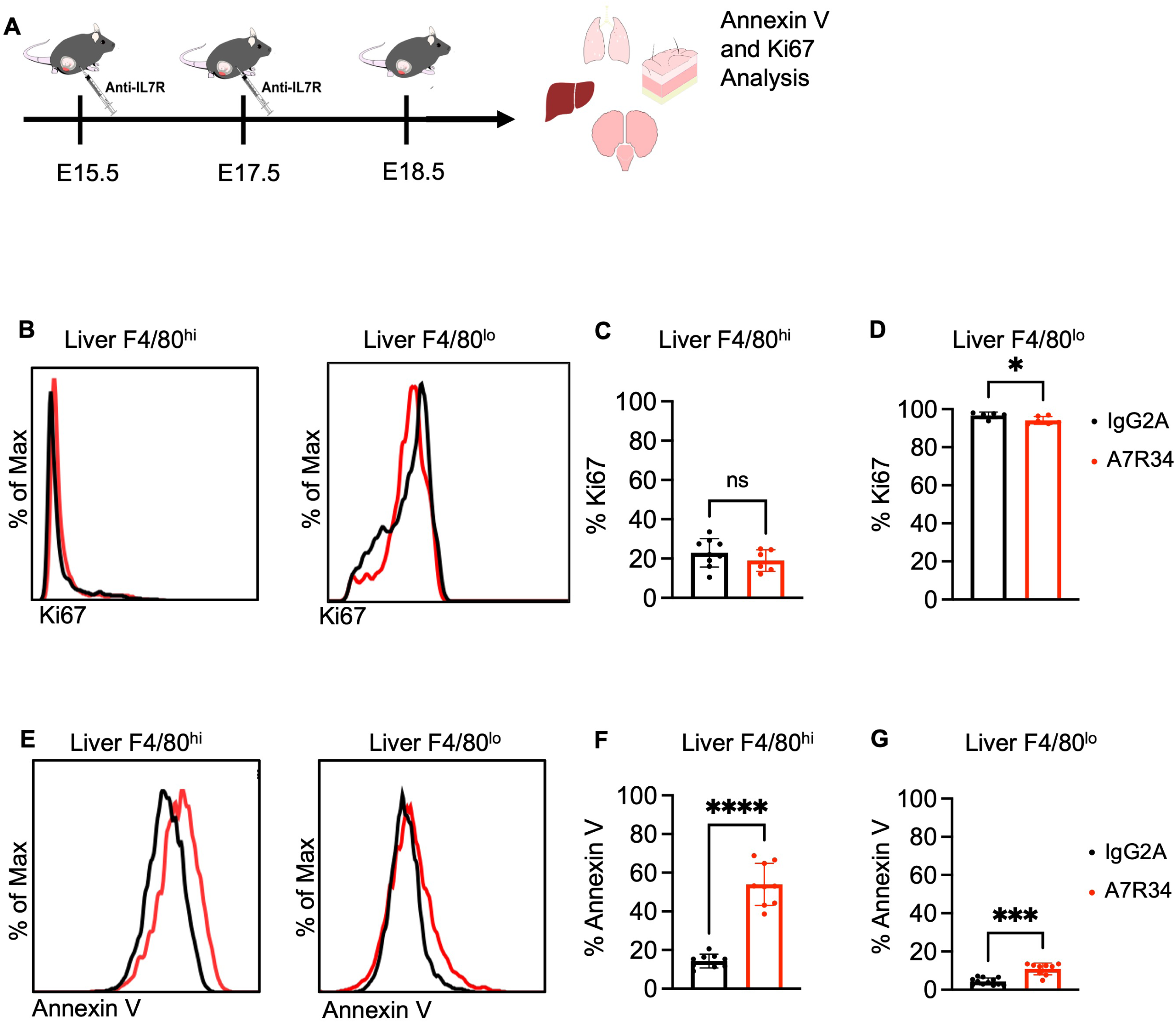
IL7/IL7R regulates proliferation and survival in fetal liver myeloid cells. **(A)** Pregnant WT dams were treated with anti-IL7R blocking antibody at embryonic days (E)15.5 and E17.5. At E18.5, myeloid populations in fetal liver were analyzed for proliferation and apoptosis. **(B-D)** Representative plots of Ki67 expression (**B**) and quantification of %Ki67+ cells in F4/80^hi^ (**C**) and F4/80^lo^ (**D**) fetal liver populations. (**E-G**) Representative plots for Annexin V expression (**E**) and quantification of Annexin V+ F4/80^hi^ (**F**) and F4/80^lo^ cells (**G**) in the fetal liver at E18.5. Data represent average ± SD for n=5-10 mice per condition representing 3-4 independent experiments. Significance determined using Mann-Whitney test, *\*p<0.05, **p<0.01, ***p<0.001, ****p<0.0001*.

### Cre-specific deletion reveals critical developmental window for IL7R expression

We previously showed that neonatal depletion of resident tissue macrophage (RTM) compartments—whether induced by prenatal IL7R blockade or by germline IL7R deletion—is followed by restoration of adult RTM cellularity^6^. This recovery might reflect either regeneration of IL7R-dependent macrophages or replacement by IL7R-independent populations. To distinguish between these possibilities, we combined inducible deletion and fate-mapping approaches to remove Il7r specifically from erythromyeloid progenitor (EMP)–derived macrophages or from hemogenic endothelium/definitive hematopoiesis–derived precursors, while simultaneously tracking Il7r-deleted cells via tdTomato expression. To achieve this, we crossed Csf1r-Mer-Cre-Mer or Cdh5-Cre-ERT2 mice to Il7r^fl/fl;Ai^^9^ reporter mice and induced Cre recombination with 4-hydroxytamoxifen at E8.5 or E10.5, respectively (**Fig. 5A**). Efficient Il7r deletion in tdTomato⁺ cells was confirmed by PCR on sorted populations (**SFig. 5A**). As expected, Csf1r-Mer-Cre-Mer pulsed at E8.5 selectively labeled microglia but not peripheral blood (PB) CD45⁺ cells (**SFig. 5B**), whereas Cdh5-Cre-ERT2 pulsed at E10.5 labeled PB CD45⁺ cells but not microglia (**SFig. 5C**), validating the fidelity of both lineage-tracing and deletion strategies.

**Figure 5.**
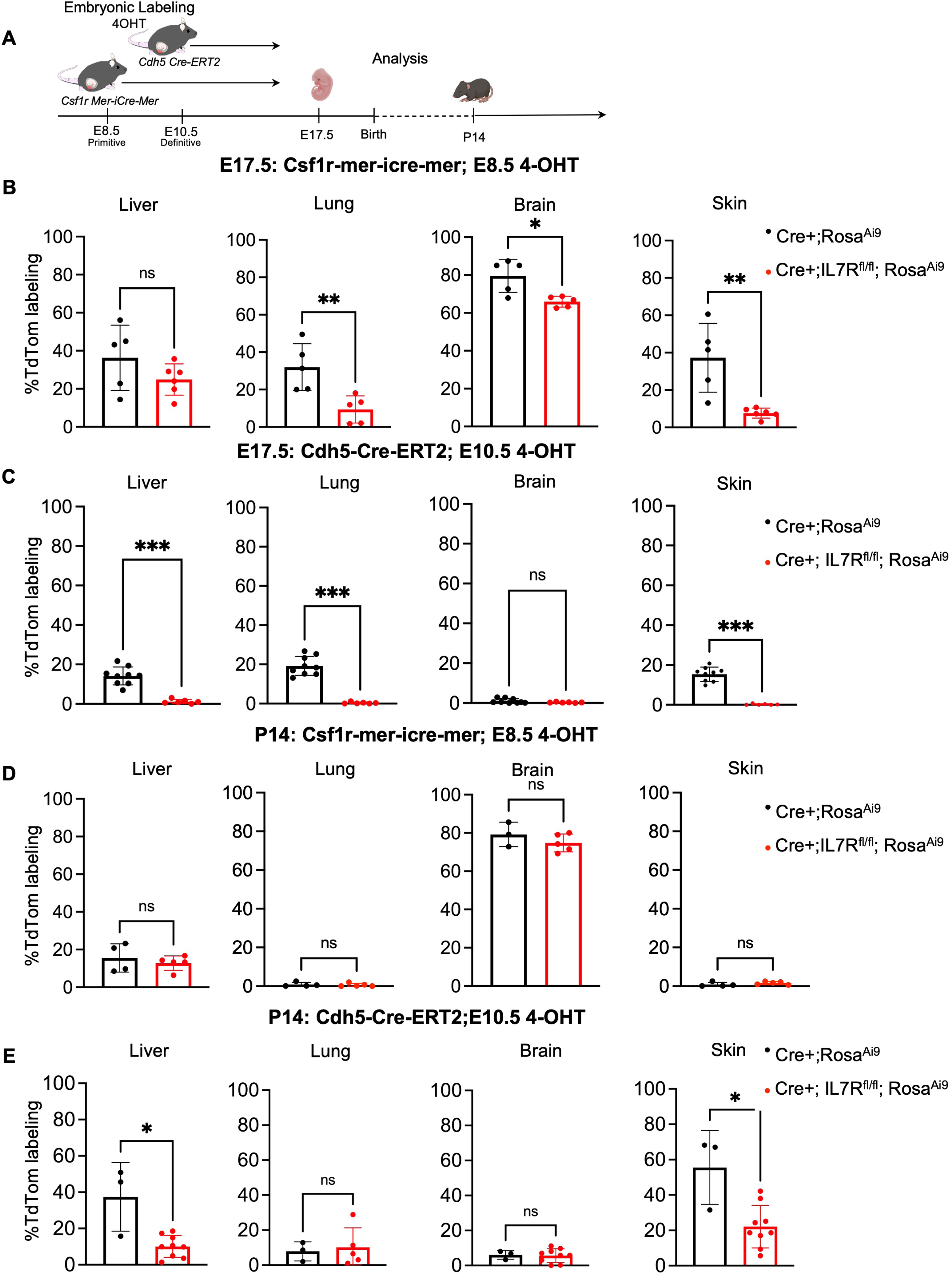
Cre-specific deletion reveals critical developmental window for IL7R expression. **A)** Schematic of fate mapping and deletion of IL7R-expressing RTMs derived from EMP and hemogenic endothelium/definitive HSC utilizing Csf1r-Mer-cre-Mer and Cdh5-Cre-ERT2 pulsed with 4-hydroxytamoxifen at E8.5 and E10.5, respectively. **B)** TdTomato labeling tracking Il7r deletion in macrophages in liver, lung, brain, and skin at E17.5 in Csf1r-Mer-cre-Mer x IL7R^fl/fl;Ai9^ embryos or Csf1r-Mer-cre-Mer x Ai9 controls. **C)** tdTomato labeling tracking Il7r deletion in macrophages in liver, lung, brain, and skin at E17.5 in Cdh5-Cre-ERT2 x IL7R^fl/fl;Ai9^ embryos or Cdh5-Cre-ERT2 x Ai9 controls. **D)** tdTomato labeling tracking IL7R deletion in macrophages in liver, lung, brain, and skin at postnatal day 14 (P14) in Csf1r-Mer-cre-Mer x IL7R^fl/fl;Ai9^ offspring or Csf1r-Mer-cre-Mer x Ai9 controls. **E)** tdTomato labeling tracking Il7r deletion in macrophages in liver, lung, brain, and skin at P14 of Cdh5-Cre-ERT2 x IL7R^fl/fl;Ai9^ offspring or Cdh5-Cre-ERT2 x Ai9 controls. Data represent average ± SD with n=3-9 per condition representing 3 independent experiments. Significance determined using Mann-Whitney test, *\*p<0.05, **p<0.01, ***p<0.001, ****p<0.0001*.

We first assessed the impact of Il7r deletion in EMP-derived macrophages by analyzing Csf1r-Mer-Cre-Mer–labeled cells at E17.5. Few labeled monocytes were detected at this stage, demonstrating minimal ongoing contribution of EMPs to the monocyte pool (**SFig. 5D**). In contrast, 20–40% of macrophages in the lung, liver, and skin, and the majority of microglia, were labeled, consistent with the established contribution of EMPs to microglia ^43^. Il7r deletion caused a modest but significant reduction in EMP-derived macrophages in the brain, lung, and skin at E17.5 (**Fig. 5B**).

We next examined Il7r deletion in definitive hematopoiesis–derived cells using Cdh5-Cre-ERT2 pulsed at E10.5. At E17.5, robust labeling of both Ly6c^hi^ and Ly6c^lo^ monocytes across all tissues was observed, and this labeling was completely eliminated following Il7r deletion (**SFig. 5F**). Cdh5-Cre-ERT2 also labeled ∼20% of RTMs in the lung, liver, and skin, but not microglia (**Fig. 5C**). Notably, all Cdh5-Cre-ERT2–labeled macrophages in these tissues were also eliminated by Il7r deletion (**Fig. 5C**).

IL7R expression during myelopoiesis is confined to a prenatal developmental window ^6^, consistent with the restriction of IL7R expression to lymphopoiesis in adulthood^11^. To determine how Il7r deletion during this window affects long-term persistence of macrophage subsets, we tracked EMP– and fetal definitive-derived macrophages into postnatal life following specific Il7r deletion (**Fig. 5D–E**). At P14, labeling of EMP contributions to microglia remained unchanged by Il7r deletion, as did labeling of RTMs in the liver, despite a transient reduction in labeling in response to prenatal Il7r deletion. By contrast, EMP contribution to lung and epidermal RTMs was minimal at P14, reflected by the low frequency of Csf1r-Mer-Cre-Mer–labeled cells at this stage (**Fig. 5D**). Strikingly, deletion of Il7r resulted in a sustained impairment in the contribution of definitive hematopoiesis–derived RTMs to the liver and epidermis at P14 (**Fig. 5E**). Because IL7R expression is absent from liver macrophages in postnatal life, these findings indicate that IL7R-expressing definitive fetal liver–derived RTMs are not replaced after the prenatal period.

### Prenatal blockade of IL7R impairs the adaptive response in a model of chronic liver fibrosis

Our data above demonstrate that IL7R expression in fetal macrophages regulated the expression of prototypical inflammatory pathways and tissue remodeling in developing myeloid cells within the fetal liver. It is well established that proinflammatory hepatic macrophages promote liver fibrosis by releasing inflammatory cues including TNF-α, IL-1β, IL-6, and TGF-β necessary for the activation of hepatic stellate cells, the main effector cell responsible for the production of extracellular matrix (ECM) in hepatic fibrosis ^44^. To test whether this developmental transcriptional phenotype has a sustained effect on tissue homeostasis, we blocked IL7R signaling by prenatal administration of anti-IL7R antibody during fetal macrophage development (**Fig. 6A**). We then subjected adult offspring to six weeks of chronic carbon tetrachloride (CCl_4_)-mediated liver fibrosis challenge beginning at 6 weeks of age. We reasoned that developmental blockade of IL7R signaling during fetal development would influence the adaptive response to chronic fibrotic liver injury, thereby promoting fibrosis in response to injury. Compared to IgG2A-treated controls, offspring treated prenatally with A7R34 IL7R blocking antibody had enhanced liver fibrosis, as evidenced by both hematoxylin and eosin staining and Masson-trichrome staining for increased deposition of collagen (**Fig. 6B**). These pathological changes were also associated with impaired liver function, reflected as increased levels of liver enzymes alanine transaminase (ALT) (**Fig. 6C**) and aspartate transaminase (AST) (**Fig. 6D**) in response to prenatal IL7R blockade.

**Figure 6.**
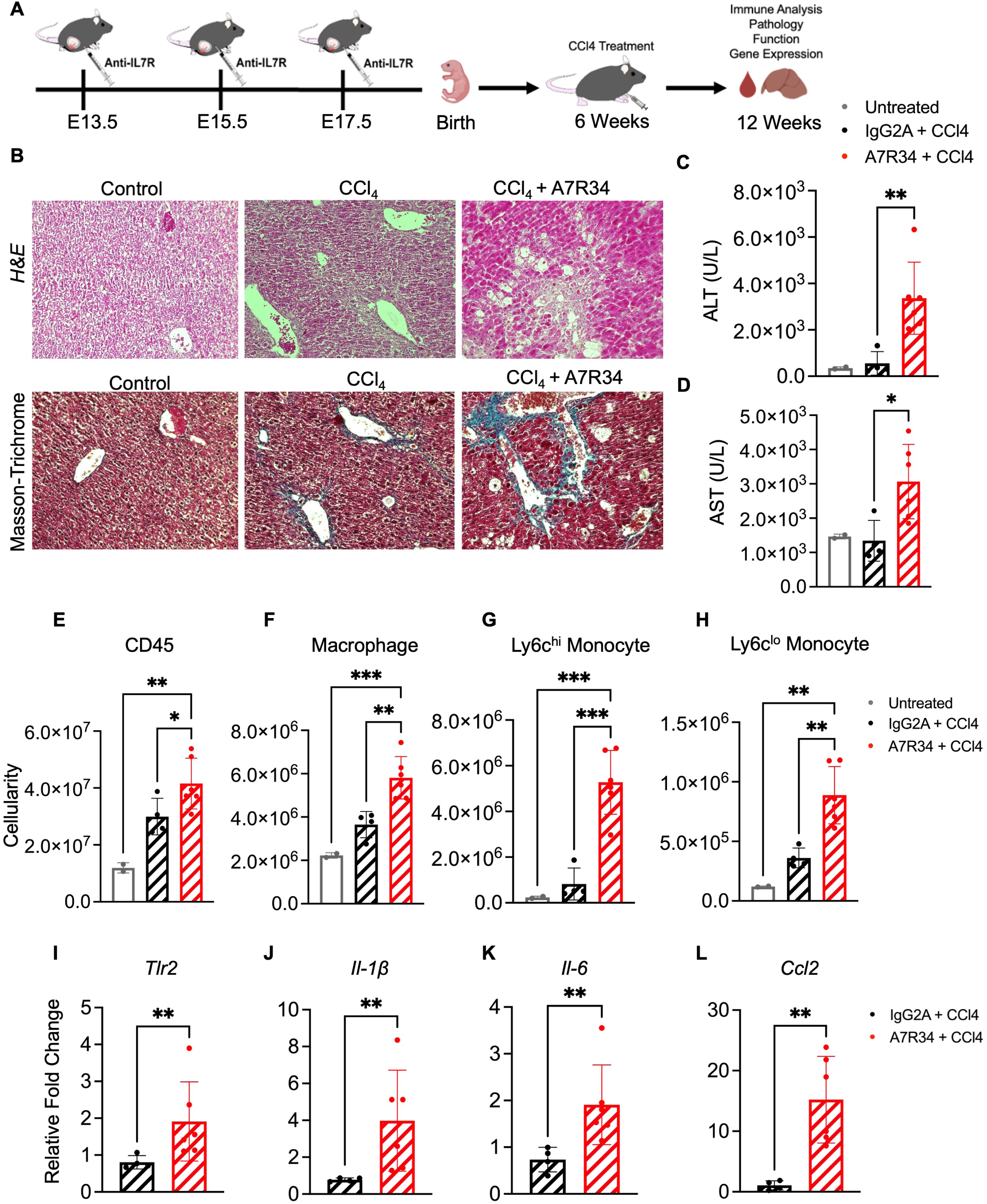
Prenatal IL7R blockade impairs the adaptive response to chronic liver injury in adulthood. **A)** Pregnant mice were treated with an IL7R blocking antibody (A7R34) or isotype antibody control during late gestation and adult female offspring were treated with CCl4(2 μl/g) 3 times a week for 6 weeks and liver fibrosis was assayed by immune analysis, pathology, function, and gene expression in the liver. **B**) Hematoxylin and Eosin (H&E) staining and Masson-Trichome staining of control (No CCl4), CCl4– and A7R43+ CCl4-treated adult liver. **C-D**) Measurement of liver enzymes alanine transaminase (ALT) (**C**) and aspartate transaminase (AST) (**D**) following chronic CCl4 treatment. **E-H**). Total cellularity of CD45 (**E**), F4/80^hi^CD11b^lo^ RTMs (**F**), F4/80^hi^CD11b^lo^Ly6c^hi^ (**G**), and F4/80^hi^CD11b^lo^Ly6c^lo^ (**H**) cells in the liver following chronic CCl4 treatment. **(I-L)** Log-fold change in gene expression of TLR2 (**I**), IL-1B (**J**), IL-6 (**K**), and CCL2 (**L**). Gene expression analysis was performed in triplicate on at least three independent samples. Data represents average ± SD of n=2-4 mice per condition representing two independent experiments. Significance determined using Mann-Whitney test, *\*p<0.05, **p<0.01*.

Prenatal IL7R blockade significantly increased both total CD45+ cellularity (**Fig. 6E**) and RTM cellularity (**Fig. 6F**) relative to IgG2A-treated controls in response to chronic CCl_4_ challenge. Increased RTM cellularity was accompanied by an increase in both Ly6c^hi^ and Ly6c^lo^ infiltrating monocytes (**Fig. 6G-H**). Neutrophils were also significantly increased **(SFig. 6A)**, but B– and T-cells numbers were significantly decreased in response to prenatal IL7R blockade **(SFig 6B-C)**. Lastly, prenatal IL7R blockade promoted a typical macrophage-mediated inflammatory response, including increased transcription of *Tlr2*, *Il-1β*, *Il-6*, *Ccl2*, *Tlr4*, and *Ccr2* (**Fig. 6I-L; SFig. 6D-E**). Expression of *Pdgfra*, a marker of hepatic stellate cell activation, was also elevated. (**SFig. 6F**). Together, these data implicate an impaired adaptive response to injury mediated by pro-inflammatory macrophages and monocyte infiltration with subsequent induction of hepatic fibrosis in adult mice after prenatal IL7R blockade.

## Discussion

Our investigation revealed that IL7 acts synergistically with its cognate receptor IL7R to regulate fetal myelopoiesis, including RTM proliferation, survival, and acquisition of anti-inflammatory functions during fetal development. IL7 was initially identified as a growth factor regulating B-cell development ^7,33^, and IL7 signaling has been studied almost entirely in the context of lymphopoiesis. We first described a role for IL7R expression in regulating prenatal RTM development, and here we build on those findings by demonstrating the functional role of the cognate ligand, IL7, during fetal myelopoiesis. Although IL7 is neither necessary nor sufficient for macrophage generation, our *ex vivo* cell culture studies demonstrated that IL7 directly impacted macrophage generation from fetal myeloid progenitors. Our analysis further revealed that IL7/IL7R signaling is co-opted for fetal myeloid lineage development to promote survival and restrain inflammatory signaling in fetal myeloid cells that seed tissues during the prenatal window.

IL7/IL7R signaling in lymphoid cells is typically characterized by rapid phosphorylation of Stat5 ^45,46^, whereas prolonged exposure results in AKT phosphorylation to promote survival and cell cycle progression via *Bcl2* upregulation and negative regulation of *p27kip*, *GSK3*, and *Bad* ^47–53^. One other report also recently described pI3K/AKT signaling in IL7R-expressing monocytes in a pathological inflammatory context--rheumatoid arthritis ^54^. Consistent with phospho-AKT signaling promoting survival in both myeloid and lymphoid cells, our transcriptional analysis indicates that IL7R signaling promotes survival and proliferation modules in fetal myeloid cells, data that is supported by our *in vivo* analysis in response to IL7R blockade. Synergistic signaling with other niche factors, such as M-CSF, likely promotes survival and polarization during niche education. Other signals also act synergistically with M-CSF to further enhance polarization and niche education, including IL-34 in microglia and Langerhans cells, and TGF-β in alveolar macrophages ^22,25^. Several studies have described the combinatorial effect of IL7R signaling and TCR signaling for an amplified TCR response in T-cells ^55,56^. A similar phenomenon may occur in fetal myeloid cells, where IL7R signaling may function to amplify other signals to regulate the acquisition of tissue-specific function.

Most RTM populations undergo “layered immune development”, a process in which early fetal-derived macrophages are gradually replaced by precursors from later developmental waves. The timing, extent, and source of this replacement are dependent on the tissue. In Il7^GFP/GFP^ and Il7r^⁻/⁻^ models, as well as in response to prenatal IL7R blockade, impaired IL7R signaling causes perinatal depletion of RTMs in peripheral tissues such as the liver. Cellularity is normalized by adulthood ^6^, presumably by other, non-IL7R-dependent sources. Our inducible deletion experiments revealed that IL7R-deficient macrophages of EMP origin were transiently lost during fetal liver development but recovered postnatally, whereas IL7R-deficient macrophages arising from hemogenic endothelium are lost in utero and fail to be replenished postnatally. Although Cdh5-Cre-ERT2 marks definitive HSCs emerging from the hemogenic endothelium, recent evidence suggests it may also mark other HSC-independent lineages whose contributions to fetal myelopoiesis and RTM population have not been investigated ^57^. These data suggest that despite normalization of postnatal RTM cellularity, loss of IL7R-dependent RTMs during fetal development is not recoverable and may have functional implications.

We tested the functional implications of loss of fetal IL7R-dependent macrophages on liver function by assessing the effect of developmental IL7R receptor blockade on induced liver fibrosis, in which we observed an impaired response to adaptive liver injury in adulthood. During the same prenatal window, developmental expression of IL7R serves as a mechanism by which fetal-derived RTMs acquire specific functions that promote tissue homeostasis. Gene expression analysis of *Il7r*-deficient myeloid cells revealed that IL7R expression regulates inflammatory gene programs during development, including TLR signaling, IL6/Jak/Stat signaling and interferon signaling. We propose that IL7R not only regulates survival of developing macrophages during fetal development, but also endows developing RTMs with an anti-inflammatory transcriptional phenotype that regulates tissue homeostasis and adaptive responses to injury. Indeed, prenatal IL7R blockade not only sensitized adult offspring to liver fibrosis in response to chronic CCl_4_-mediated liver injury, increased RTM cellularity and monocyte infiltration in response to CCl_4_-mediated liver injury, and was associated with higher levels of inflammation, including TLR signaling, IL-6 and IL-1β expression, suggesting that replacement of IL7R-expressing fetal RTMs can be deleterious. Fetal IL7R blockade also likely impaired lymphoid development, although we allowed ample time for recovery before challenge, and we previously demonstrated normalization of the lymphoid compartment in adulthood. However, it is possible that the development of specific developmentally-restricted *γ*δ-T and T-regulatory cells was impacted in our model. These subsets have been implicated in mediating liver fibrosis in prior studies ^58,59^; however, mechanistic work to examine these associations has relied on broader deletion strategies targeting γδ T cells or Treg compartments as a whole ^60,61^. Therefore, the relative contribution of developmentally restricted subsets to liver fibrosis outcomes remains unclear and warrants further investigation. Together, these data suggest that IL7R expression in fetal macrophages has a functional effect on the adaptive response to injury in adulthood, and that replenishment of the RTM compartment –– in the absence of IL7R expression--does not functionally replicate the role of IL7R-dependent macrophages in adult tissues. Our data provide an additional line of evidence that fetal-derived RTMs have specific homeostatic functions in adult tissues and implicate developmental IL7R signaling in conferring anti-inflammatory properties and adaptive response to injury.

## Acknowledgements

We thank the Dr. K. Ikuta for the IL7-GFP mice; Dr. David Gravano of the UC Merced Stem Cell Instrumentation Foundry and James Marvin of the University of Utah Flow Cytometry Core for their flow cytometry support; Drs. Shannon Buckley, Diego López, and Hans Haecker for helpful comments on the manuscript. This work is supported by NIH/NHLBI R01HL147081 to AEB. AEB is also the recipient of NIH/NHLBI award K01HL130753, the Pew Foundation Biomedical Scholars Award, and the Hellman Fellows Award from the Hellman Foundation. GAL was supported by T32DK007115, NAT is supported by NIH F31HL182256, BPK is supported by NIH F31HL170732, and CHV is supported by T32AI138945. PNM is supported by NIH R21 DK115991 and NIH R01 DK128819.

## Author Contributions

AEB was responsible for supervision, conceptualization, experimental design, manuscript preparation, and funding. GAL was responsible for experimental design, data collection, data analysis, data visualization, computational analysis, and manuscript preparation. NAT was responsible for data collection, data analysis, data visualization, computational analysis, and manuscript revision. BPK was responsible for image collection, image analysis, and figure preparation. CHV, SA, NH, DAB, and EMK were responsible for data collection and data visualization. PNM was responsible for experimental design and manuscript revision.

## Conflict of interest Disclosure

The authors declare no conflicts of interest.

## Notes

### Competing Interest Statement

The authors have declared no competing interest.

