## Supplemental Materials for "Developmental IL7/IL7R signaling in fetal resident tissue macrophages regulates adaptive responses to liver injury"

**Supplementary Data**

**Supplementary Figure 1. IL-7 expression and function during fetal myeloid development. A)** Frequency of neonatal (P4) B-cells within total CD45+ cells in *Il7* $^{GFP/GFP}$ mice compared to WT. Data represents average ± SD for n=9 mice per condition representing three independent experiments. **B,C)** Frequency of adult (8-10 weeks) B- cell (**B**) and T-cells (**C**) within total CD45+cells in *Il7*$^{GFP/GFP}$ mice compared to WT. Data represents average ± SD of n=6-12 mice per condition representing four independent experiments. Significance determined using Mann-Whitney test, **p<0.05, **p<0.01.* **D-F)** Immunofluorescence staining of IL7R+ F4/80+ cells in close proximity to IL7 expression in (**D**) lung, (**E**) brain**,** and (**F**) skin of E14.5 murine embryos. Slides were imaged on a Zeiss LSM 700 Zen Black software and 63x objective, images generated using ImageJ software. Rectangular fields show images at magnification and highlight IL7R+ F4/80+ cells colocalized with IL7+ staining. IL7R (red) indicated by arrowhead, F4/80 (green) indicated by arrow, and IL7 (gray) indicated by fishtail arrow. Images are representative of three individual embryos.

**Supplementary Figure 2. IL-7 expression regulates fetal myeloid development.**

**A,B)** Representative FACS plots of myeloid cells in the liver, lung, brain, and epidermis in the neonate (P4, **A**) and adult (8-10 weeks, **B**) in $IL7^{GFP/GFP}$ mice and WT controls. C) Cellularity of $F4/80^{hi}$ macrophages generated after 6 days of culture of E14.5 WT fetal MDPs, fetal cMOPs and fetal MPs cultured in in m-CSF (20 ng/ml) or rIL7(100 ng/ml) alone. Data represents average ± SD from three individual experiments and progenitors plated in triplicate. Significance determined by paired t-test of averaged triplicate values, **p<0.05, **p<0.01, ***p<0.001.*

**Supplementary Figure 3. Regulating of downstream signaling in fetal myeloid cells by IL7R signaling.**

**A**) Representative gating of sort strategy for RNA-seq experiments in **Figures 2 and 3**. **(B-G)** Representative histograms and accompanying quantification of phospho-Stat5 signal in F4/80$^{hi}$ and F4/80$^{lo}$ fetal liver cells after stimulation with rGM-CSF (50 ng/mL) **(B,C)**,rIL7 (100 ng/ml) **(D,E)**, and rGM-CSF + rIL7 **(E,F)** for 5 minutes. Data are representative of three individual experiments. Significance determined using Mann-Whitney test, *$p<0.05$.*

**Supplementary Figure 4. Prenatal IL7R blockade impairs proliferation and survival in fetal myeloid cells. (A)** Analysis and quantification of Ki67 expression in F4/80$^{hi}$ and F4/80$^{lo}$ cells in lung, brain, and skin at E18.5 following prenatal IL7R blockade (see **Fig. 4A**). **(B)**. Analysis and quantification of Annexin V staining in F4/80$^{hi}$ and F4/80$^{lo}$ cells in lung, brain, and skin at E18.5 following prenatal IL7R blockade. Data represent average ± SD with n=5-10 per condition representing 3 independent conditions. Significance determined using Mann-Whitney test, **p<0.05, **p<0.01, ***p<0.001, ****p<0.0001*.

**Supplementary Figure 5. Inducible Cre-specific deletion of Il7r. A)** Gel electrophoresis of PCR products from Tomato+ and Tomato- cells from Cdh5-Cre-ERT2 x Ai9 mice. The gel displays PCR products at their expected lengths, with IL7R WT cells (Tom-) at 325 bp and IL7R mutant (Tom+) at 162 bp. IL7R deletion is confirmed in tdTomato+ cells in the Cdh5-Cre fate mapping model and IL7R WT is present in tdTomato- cells*.* **B)** %TdTomato labeling in Csf1r-mer-Cre-mer X Ai9 mice induced by injection of 4-OHT at E8.5 in microglia and peripheral blood determined at P0. C) %TdTomato labeling in Cdh5-Cre-ERT2XAi9 mice induced by injection of 4-OHT at E10.5 in microglia and peripheral blood determined at P0. **D**) TdTomato labeling of Ly6c$^{hi}$ and Ly6c$^{lo}$ monocytes in brain, liver, lung and skin of Embryonic Day (E) 17.5 Csf1r-mer-Cre-mer X IL7R$^{fl/fl}$;Ai9 or Ai9 embryos. **D)** TdTomato labeling of ly6c$^{hi}$ and ly6c$^{lo}$ monocytes in brain, liver, lung and skin of Embryonic Day (E) 17.5 Cdh5-Cre-ERT2X IL7R$^{fl/fl}$;Ai9 or Ai9 embryos.

**Supplementary Figure 6. Effects of prenatal IL7R blockade in a CCl4-mediated chronic liver injury model. (A-C).** Total cellularity of neutrophils (**A**), B Cells (**B**) and T Cells (**C**) in the liver following chronic $CCl_4$ treatment. **(D-F)** Log-fold change in gene expression of TLR4 (**D**), CCR2 (**E**), and PDGFRα (**F**). Gene expression analysis was performed in triplicate on at least three independent samples. Data represents average ± SD of n=2-4 mice per condition representing two independent experiments. Significance determined using Mann-Whitney test, **p<0.05, **p<0.01.*

**SFig 1. IL7 expression and function during fetal myeloid development.**

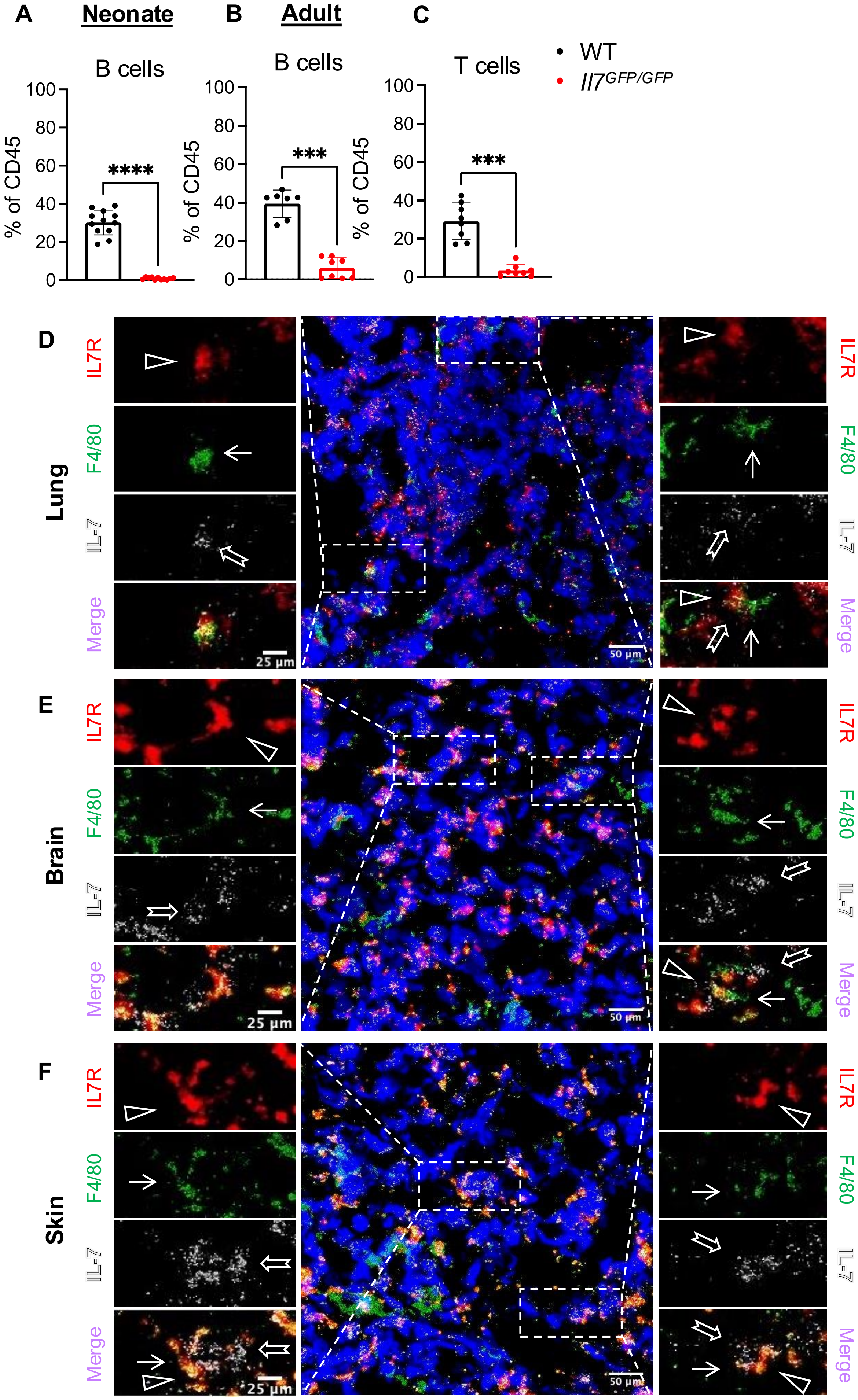

**SFig 2. IL-7 expression regulates fetal myeloid development.**

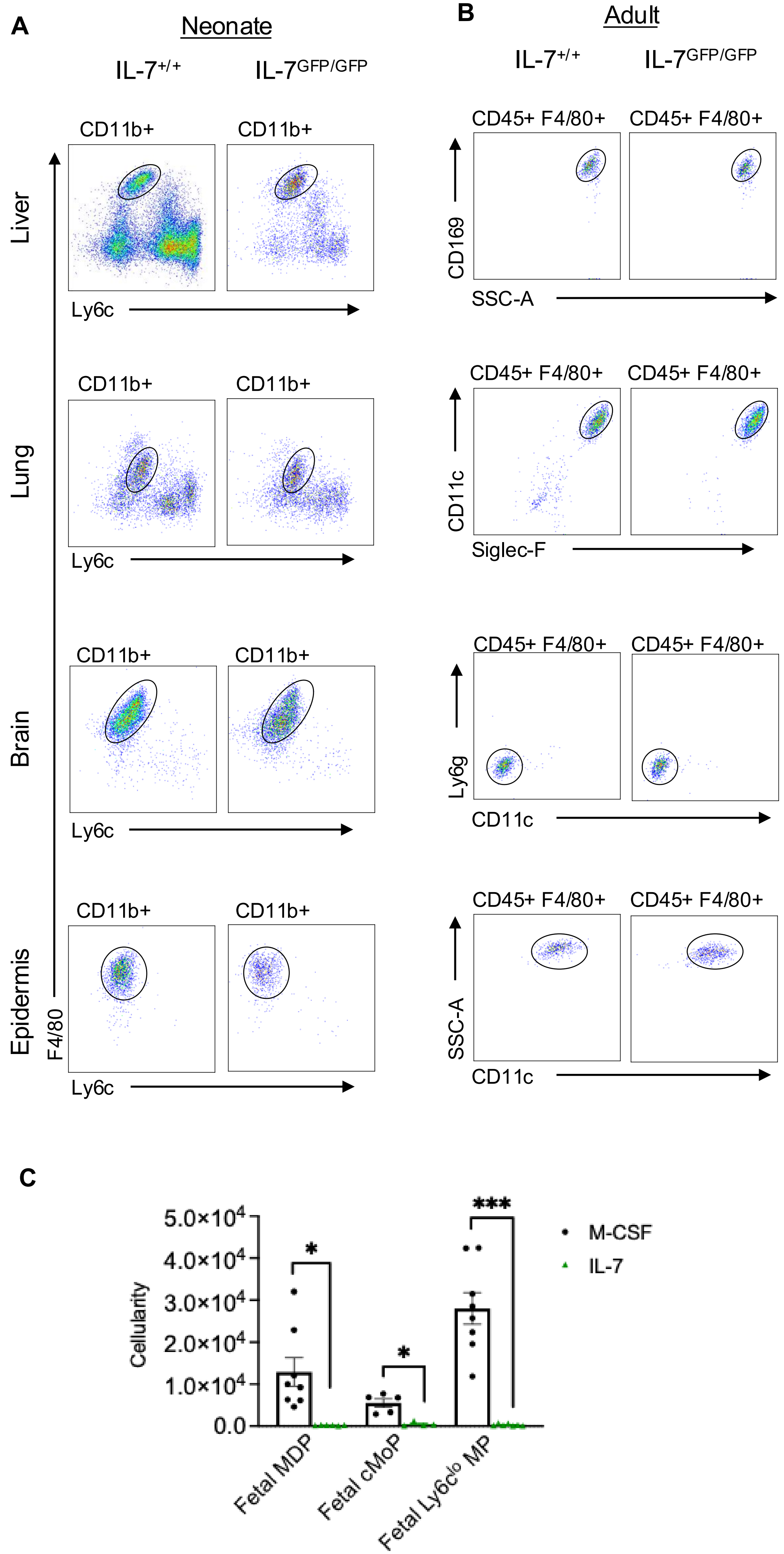

**SFig 3. Regulating of downstream signaling in myeloid cells by IL7R signaling.**

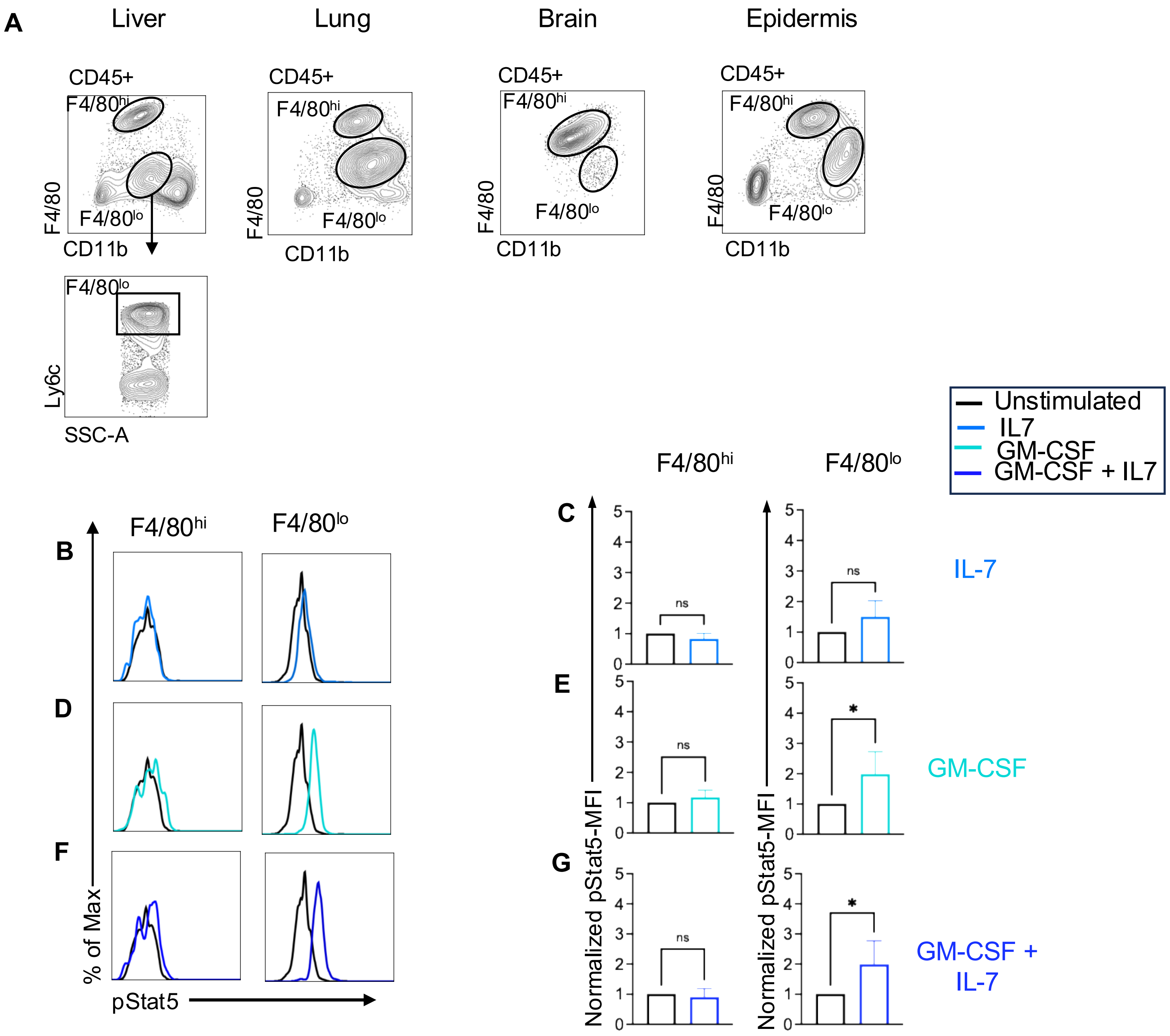

**SFig 4. Prenatal IL7R blockade impairs proliferation and survival in fetal myeloid cells.**

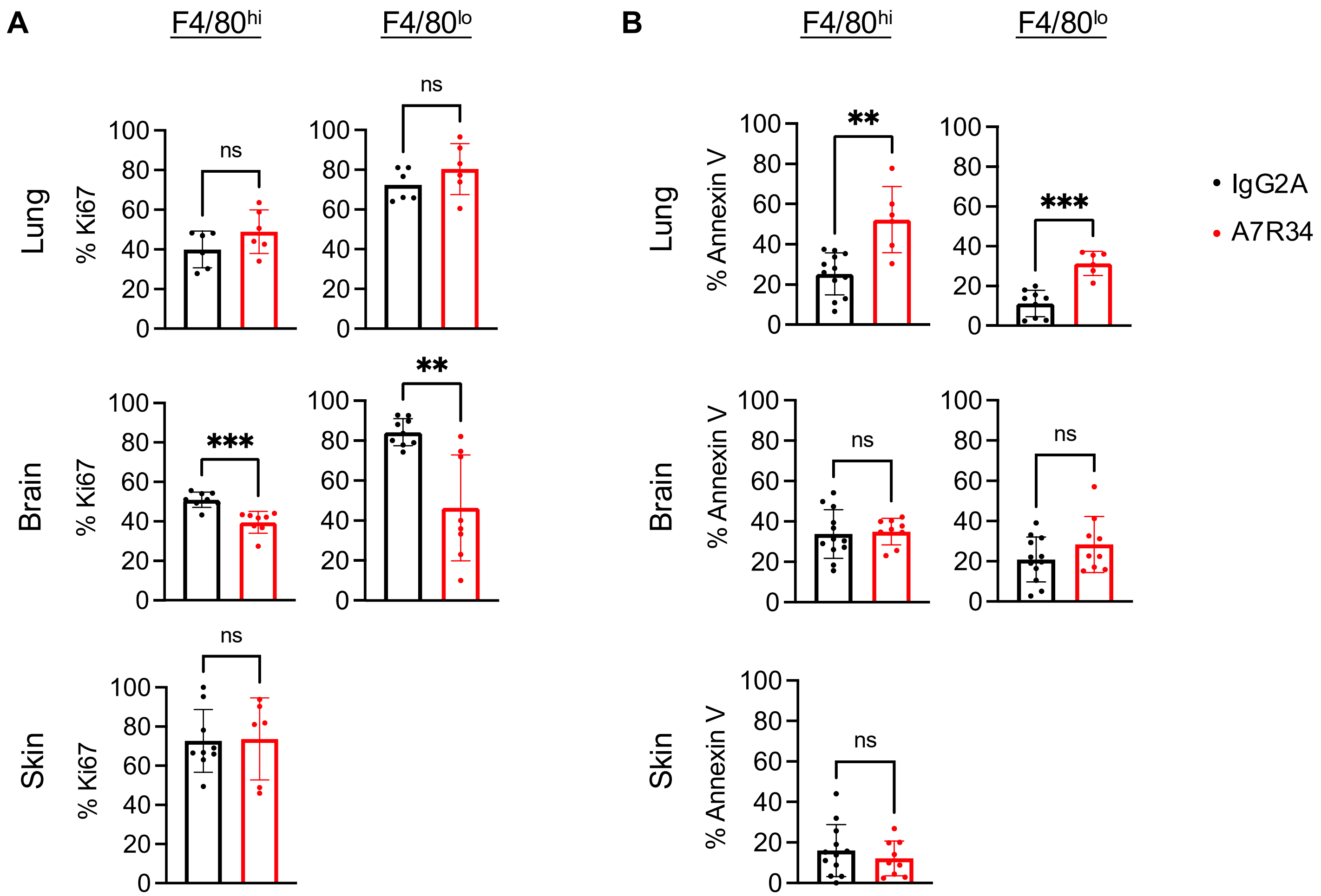

**SFig 5. Specific deletion of IL7R impacts fetal definitive contribution to monocytes**

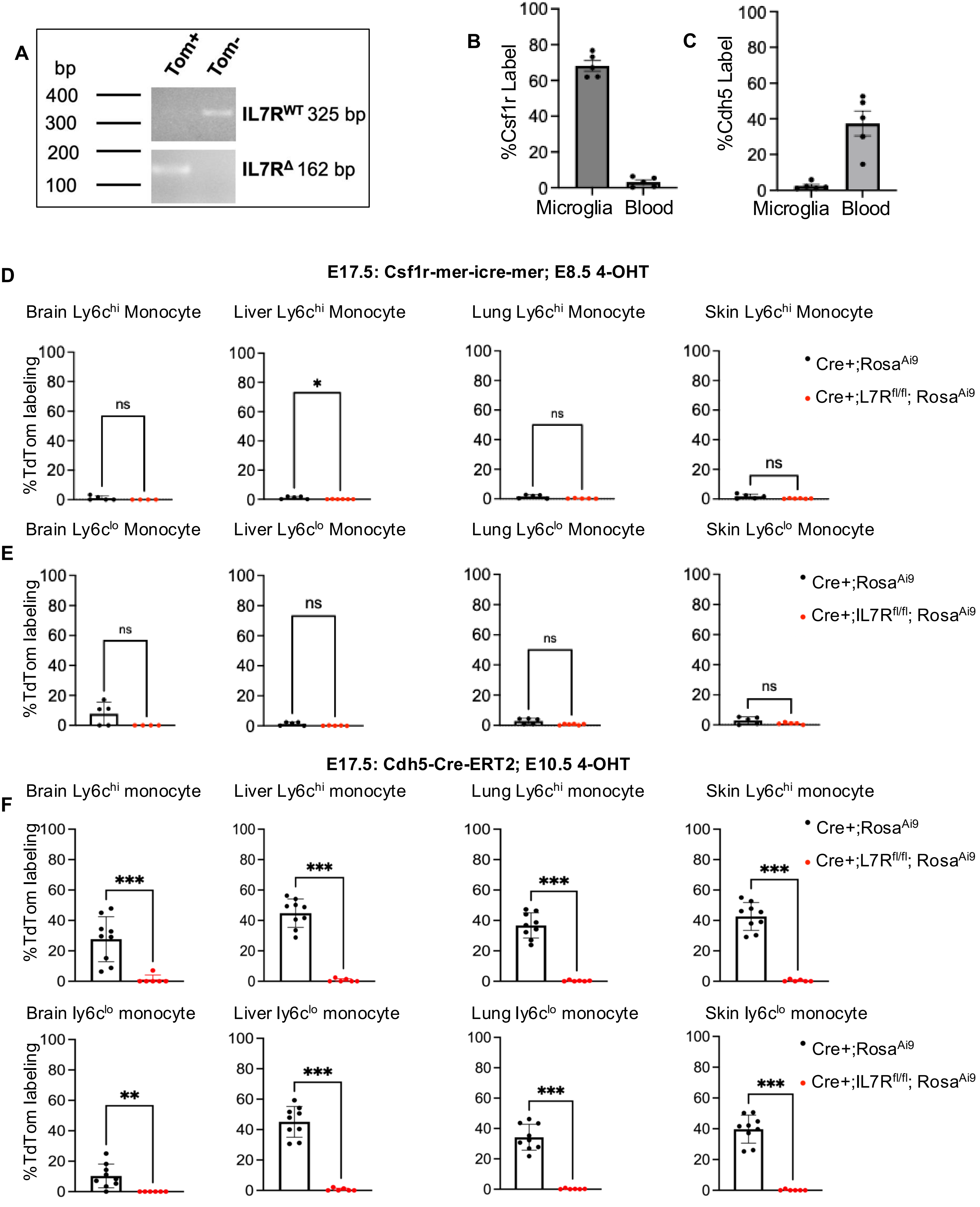

**SFig 6. Effects of prenatal IL7R blockade in a CCl4-mediated chronic livery injury model**

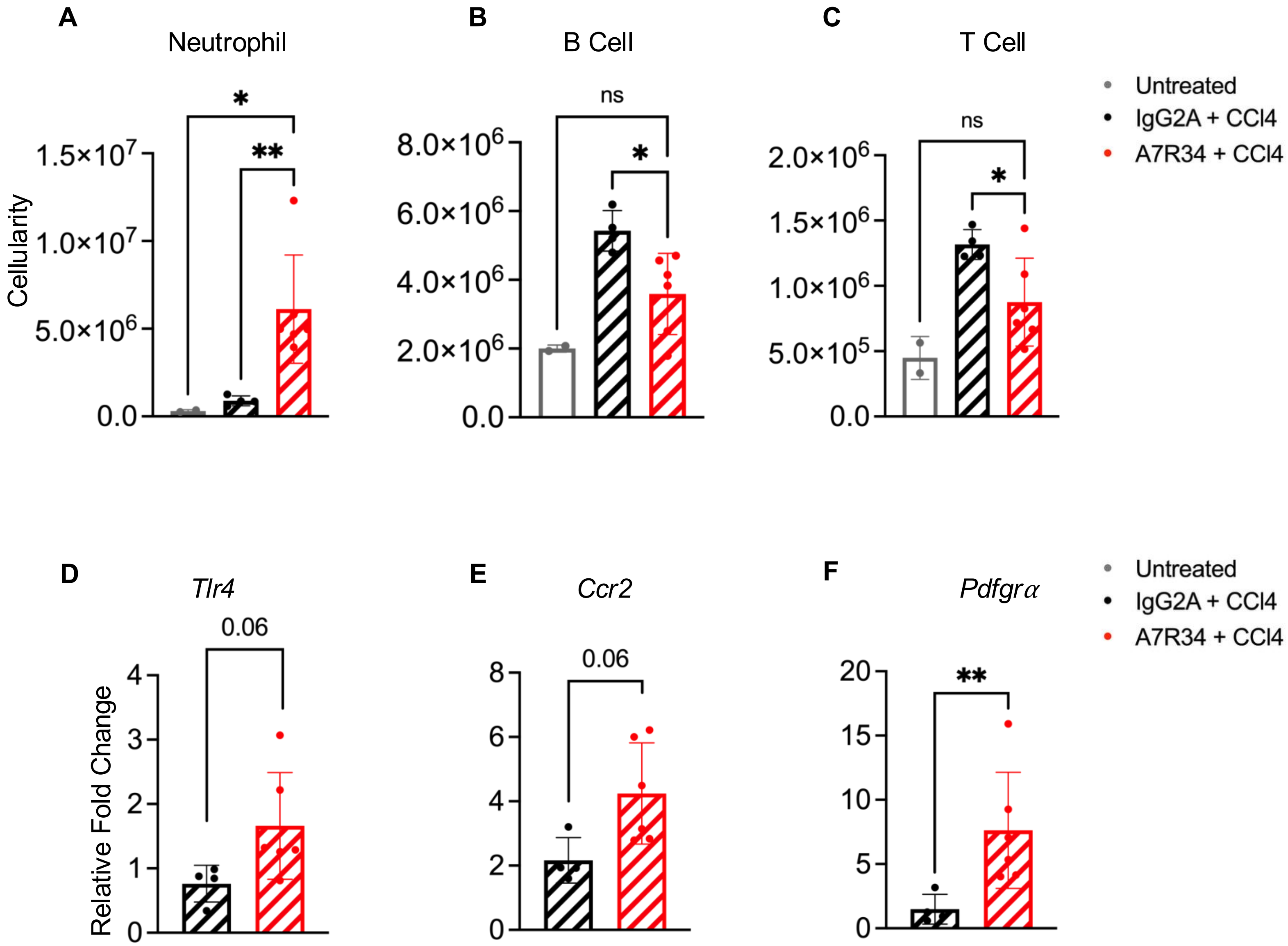
